# Emotional prosody in congenital amusia: impaired and spared processes

**DOI:** 10.1101/466748

**Authors:** A. Pralus, L. Fornoni, R. Bouet, M. Gomot, A. Bhatara, B. Tillmann, A. Caclin

## Abstract

Congenital amusia is a lifelong deficit of music processing, in particular of pitch processing. Most research investigating this neurodevelopmental disorder has focused on music perception, but pitch also has a critical role for intentional and emotional prosody in speech. Two previous studies investigating amusics’ emotional prosody recognition have shown either some deficit or no deficit (compared to controls). However, these previous studies have used only long sentence stimuli, which allow for limited control over acoustic content. Here, we tested amusic individuals for emotional prosody perception in sentences and vowels. For each type of material, participants performed an emotion categorization task, followed by intensity ratings of the recognized emotion. Compared to controls, amusic individuals had similar recognition of emotion in sentences, but poorer performance in vowels, especially when distinguishing sad and neutral stimuli. These lower performances in amusics were linked with difficulties in processing pitch and spectro-temporal parameters of the vowel stimuli. For emotion intensity, neither sentence nor vowel ratings differed between participant groups, suggesting preserved implicit processing of emotional prosody in amusia. These findings can be integrated into previous data showing preserved implicit processing of pitch and emotion in amusia alongside deficits in explicit recognition tasks. They are thus further supporting the hypothesis of impaired conscious analysis of pitch and timbre in this neurodevelopmental disorder.

**Highlights:**

- Amusics showed preserved emotional prosody recognition in sentences
- Amusics showed a deficit for emotional prosody recognition in short voice samples
- Preserved intensity ratings of emotions in amusia suggest spared implicit processes

## 1. Introduction

Congenital amusia is a lifelong deficit of music perception and production, also referred to as “tone deafness”. This disorder is estimated to affect one to four percent of the general population (Peretz, Cummings, & Dubé, 2007; Peretz & Vuvan, 2017) and is suggested to have genetic origins (Peretz et al., 2007). Amusic individuals have neither peripheral auditory deficits nor brain lesions, but they are unable to detect out-of-key notes in a melody, and they sing out-of-tune (Peretz, 2016; Tillmann, Albouy, & Caclin, 2015). Pitch processing deficits were observed for amusic individuals in perception tasks, such as pitch discrimination (Hyde & Peretz, 2004) or pitch contour change detection (Peretz, Champod, & Hyde, 2003), as well as in short-term memory related to pitch (Tillmann, Lévêque, Fornoni, Albouy, & Caclin, 2016). In a typical (non-amusic) brain, a fronto-temporal network is involved in pitch processing and memory, and thus also in music perception (Gaab, Gaser, Zaehle, Jancke, & Schlaug, 2003; Koelsch et al., 2009; Zatorre, Evans, & Meyer, 1994). In the amusic brain, anatomical and functional abnormalities have been observed in this fronto-temporal network (Albouy, Mattout, et al., 2013; Hyde et al., 2007; Hyde, Zatorre, Griffiths, Lerch, & Peretz, 2006; Hyde, Zatorre, & Peretz, 2011). More specifically, decreased fronto-temporal connectivity was observed in congenital amusia, in particular in the right hemisphere, together with an increased connectivity between the auditory cortices (Albouy, Mattout, et al., 2013; Albouy, Mattout, Sanchez, Tillmann, & Caclin, 2015; Hyde et al., 2011; Leveque et al., 2016; Loui, Alsop, & Schlaug, 2009; Tillmann et al., 2015). These findings suggest an altered auditory neural network underlying the pitch processing deficit in congenital amusia (Leveque et al., 2016). The deficit extends to timbre (Stewart, 2011; Tillmann, Schulze, & Foxton, 2009), whereas temporal processing seems to be mostly preserved in amusia (Hyde & Peretz, 2004), at least when the material does not entail pitch variations (Foxton, Nandy, & Griffiths, 2006; Pfeuty & Peretz, 2010).

Given these processing deficits for music, emotion processing has also been studied with musical material in congenital amusia. Despite impaired perception and memory of music, some listeners afflicted with congenital amusia have been reported to either like or avoid listening to music. This dichotomy occurs independently of the severity of amusia as measured by the Montreal Battery for the Evaluation of Amusia (Mcdonald & Stewart, 2008; Omigie, Müllensiefen, & Stewart, 2012). These subjective reports about liking/avoidance inspired recent studies investigating musical emotion processing in congenital amusia. Gosselin et al (2015) showed no impairment of emotion recognition (Gosselin et al., 2015), while Leveque et al. found a mild impairment (Lévêque et al., 2018). Similarly, a study that focused on dissonance/consonance judgments of musical materials reported that congenital amusics were able to recognize the suggested musical emotions, but they based their judgments more on roughness rather than on the harmonicity cues used by control participants (Marin, Thompson, Gingras, & Stewart, 2015). These findings and previous reports of the perceptual deficits in amusia suggests that amusics’ emotional judgments in music are based largely on roughness and tempo rather than harmonicity and mode cues (Gosselin et al., 2015; Lévêque et al., 2018).

As pitch processing is involved not only in music processing, but also in speech processing, several studies have focused on speech perception abilities in amusia. Interestingly, while early studies did not report deficits in speech processing (Ayotte, Peretz, & Hyde, 2002) or in memory for verbal materials (Tillmann et al., 2009; Williamson & Stewart, 2010), subsequent studies using more fine-grained materials and methods did reveal speech processing impairments in amusia. Specifically, intonation recognition and perception of speech contour is impaired across languages - this includes tonal languages, non-tonal languages, and even artificial verbal materials (Jiang, Hamm, Lim, Kirk, & Yang, 2010, 2012; Liu, Jiang, Francart, Chan, & Wong, 2017; Liu, Jiang, Wang, Xu, & Patel, 2015; Nan, Huang, Wang, Liu, & Dong, 2016; Nguyen, Tillmann, Gosselin, & Peretz, 2009; Tillmann, Burnham, et al., 2011; Tillmann, Rusconi, et al., 2011). In speech, prosody is essential to detect a speaker’s intentions and emotions, that is, to understand what the speaker means and to follow and participate in a conversation. Pitch changes related to prosody can be of rather large extent, and other acoustic cues (e.g., loudness, duration) also contribute to prosody. The cerebral correlates of prosody processing involve bilateral inferior frontal gyri (Frühholz, Ceravolo, & Grandjean, 2012). In addition, intentional prosody processing also involves the middle superior temporal gyrus, whereas emotional prosody processing involves the right anterior superior temporal gyrus (Frühholz et al., 2012; Liebenthal, Silbersweig, & Stern, 2016). To decode the emotional meaning of speech, the amygdala, the insula and the auditory cortex are also involved (Frühholz, Trost, & Kotz, 2016). These regions are also more generally involved in decoding emotional information, whether it occurs in music or speech materials (Bestelmeyer, Maurage, Rouger, Latinus, & Belin, 2014; Escoffier, Zhong, Schirmer, & Qiu, 2013). As the right superior temporal and inferior frontal regions participate in emotional prosody processing, and these regions exhibit differences in amusia compared to controls (Albouy, Mattout, et al., 2013; Hyde et al., 2007), Liu et al. (2015) suggested that differential processing in these regions could underlie amusics’ difficulties processing subtle emotional and prosodic changes (Ayotte et al., 2002; Patel, Foxton, & Griffiths, 2005). Thompson et al. (2012) showed a deficit in the recognition of emotional prosody in amusic individuals, especially for happiness, tenderness, irritation and sadness. They suggested that this deficit might be due to the impaired processing of acoustic features that are relevant for emotional information in music and speech, such as, for example, pitch. In contrast, Lolli et al. (2015) did not find any differences in emotion recognition in speech between amusics and controls. However, amusic individuals were impaired when speech was low-pass filtered, suggesting that low frequencies in speech carry emotional content and that amusics are less sensitive to these cues. Thompson et al. and Lolli et al. used spoken sentences of the Macquarie Battery of Emotional Prosody (MBEP; the same 84 sentences were used in both studies), which were read by actors in a natural way. In this material, numerous acoustic parameters vary depending on the emotion conveyed, so amusic participants could infer emotions using non-pitch cues, such as the intensity or the duration of vowels, words, or sentences. In addition, the differences between the results of these two studies could be due to differences in amusic recruitment criteria. Lolli et al. recruited amusics based on their pitch discrimination threshold, whereas Thompson et al. recruited amusics based on the three MBEA pitch subtests (scale, interval, contour). To further explore the potential auditory emotion processing deficit in amusia, Lima et al. (2016) investigated amusics’ emotion recognition not only in speech, but also in non-verbal auditory stimuli. Participants heard non-verbal vocalizations (e.g. screams, sobs, sighs) that were not directly related to speech and were asked to judge the degree of expressed emotion on multiple seven-point scales. Results revealed that response accuracy was decreased in congenital amusic individuals relative to the control participants for both verbal and non-verbal stimuli. However, the task involving multidimensional ratings requires making multiple comparisons, so it also involves memory. Some studies have suggested that amusia could be due to a deficit in pitch memory rather than in pitch discrimination (Albouy, Cousineau, Caclin, Tillmann, & Peretz, 2016; Albouy et al., 2015; Albouy, Schulze, Caclin, & Tillmann, 2013; Tillmann, Lévêque, et al., 2016; Tillmann et al., 2009). A memory deficit such as this could have influenced Lima et al.’s results. Lima et al. also investigated the role of acoustic parameters (F0, intensity and spectral center of gravity) in emotion recognition and showed that the combinations of acoustic cues used to determine prosodic emotions differed between amusics and controls. However, the variation in the duration of the stimuli and their rhythmic structure (specifically, laughter vs. other stimuli) across emotions could have provided a variable amount of acoustic cues in the stimuli depending on the emotion, making it difficult to pinpoint the impairment for emotion processing in amusia. The conflicting results of previous research on emotional prosody processing in congenital amusia led us in the present study to use not only sentences, but also vowels, which allowed us to reduce the variation of acoustic parameters within the stimuli. In isolated vowel sounds, temporal cues (such as duration or syllable rate differences) can be controlled, whereas this is not possible in full sentences. In addition, we used a paradigm including two judgments of the verbal stimuli; participants were first asked to recognize the emotion with a forced-choice task, and then to judge the intensity of the recognized emotion. This paradigm allowed us to evaluate whether the deficit of amusia is in recognizing an emotion (as measured with the categorization task) or occurs as part of a more general evaluation of an emotion (as measured with the intensity ratings).

Previous studies investigating emotions in music and speech materials in congenital amusia used mostly explicit emotion recognition tasks, requiring participants to choose among given categories. However, indirect investigation methods have revealed preserved implicit processing of pitch and music in amusia alongside disrupted explicit processing (Omigie, Pearce, Williamson, & Stewart, 2013; Tillmann, Albouy, Caclin, & Bigand, 2014; Tillmann, Gosselin, Bigand, & Peretz, 2012; Tillmann, Lalitte, Albouy, Caclin, & Bigand, 2016). For example, amusic individuals process pitch changes or pitch incongruity (Peretz, Brattico, Järvenpää, & Tervaniemi, 2009; Zendel, Lagrois, Robitaille, & Peretz, 2015), even if they cannot explicitly detect it (Moreau, Jolicoeur, & Peretz, 2009; Tillmann, Lévêque, et al., 2016). Based on these observations, Leveque et al. also asked participants to rate the intensity of the emotions they previously recognized in musical excerpts, as a potential measure of implicit knowledge of musical emotions (Lévêque et al., 2018). Indeed, intensity ratings of emotions can be done with less precise internal representations of these emotions. No verbal or categorical representation of the emotion is necessary; a global appreciation of the stimulus suffices (Lévêque et al., 2018). Interestingly, normal intensity ratings were observed in amusics compared to controls, suggesting that implicit processing of music-induced emotions is preserved in amusia (Lévêque et al., 2018). More generally, recent studies on congenital amusia suggest that this developmental deficit might be a disorder of consciousness and/or conscious access to pitch representations (Albouy et al., 2016; Marin et al., 2015; Moreau et al., 2009; Moreau, Jolicœur, & Peretz, 2013; Omigie et al., 2013; Peretz, 2016; Stewart, 2011; Tillmann, Lalitte, et al., 2016). Therefore, in our study, we also asked participants to rate the intensity of the just-recognized emotion as this measure could reflect implicit processing of emotional prosody in congenital amusia.

In sum, emotion perception in amusia is still understudied, in particular for speech, with conflicting results for full sentence materials. Individuals’ capacity to process prosody might depend on the type of speech stimuli used; for example, in sentences and non-verbal emotional stimuli, the amount and types of temporal cues might differ between emotions, which is not the case for short vowel stimuli. Prosody-processing capacity might also depend on the type of task used, specifically whether it is explicit or implicit. The present study tested the perception of emotional prosody in congenital amusics and matched controls with sentences and short vowels using both an explicit categorization task and intensity ratings (to access a more implicit processing of the emotion). We hypothesized that stronger deficits will be observed for emotion categorization in amusics with vowel stimuli than with sentence stimuli. Given that single vowels convey fewer temporal cues, listeners are forced to rely on pitch and other fine spectral and spectro-temporal cues to recognize emotional content. In addition, given recent findings of preserved emotional intensity judgements with musical stimuli in congenital amusia (Lévêque et al., 2018), we hypothesized that intensity ratings would also be preserved for speech stimuli, both for sentences and vowels.

## 2. Methods

### 2.1. Participants

Eighteen amusic participants and eighteen control participants matched for gender, age, laterality, education, and musical training were included in the study (**Table 1**). They all gave written informed consent to participate in the experiment. Prior to the main experiment, all participants were tested with a subjective audiometry, the Montreal Battery of Evaluation of Amusia (Peretz et al., 2003) to diagnose amusia, and a Pitch Discrimination Threshold (PDT) test (Tillmann et al., 2009). A participant was considered amusic if he/she had a global MBEA score below 23 (maximum score = 30) and/or a MBEA pitch score (average of the first three subtests of the MBEA) inferior to 22 (maximum score = 30). All control participants had a global MBEA score higher than 24.5 and a MBEA pitch score higher than 23.3 (see **Table 1**). All participants had normal hearing (hearing loss inferior to 30 dB at any frequency in both ears). Study procedures were approved by a national ethics committee and participants were paid for their participation.

**Table 1:** Characteristics of the participants in both groups. The MBEA (Montreal Battery for the Evaluation of Amusia, Peretz et al., 2003) score corresponds to the average of the six subtests of the battery (maximum score = 30, cut off: 23). Pitch mean score corresponds to the average of the three pitch subtests in the MBEA (scale, contour and interval, cut off: 22). PDT: Pitch Discrimination Threshold (see Tillmann et al., 2009). Musical emotion score corresponds to the average response to the 14 questions about musical emotion in the questionnaire (high score corresponds to high emotional reaction to music). Speech perception score corresponds to the average response to the 7 questions about speech comprehension in the questionnaire (low score corresponds to a good speech comprehension). The standard deviation is indicated in parentheses. Groups were compared with t.tests (two sided), except for handedness where a Chi2 test was used (Qobs= 0.36).

|  | Amusics (n=18) | Controls (n=18) | p-value (group comparison) |
| --- | --- | --- | --- |
| Age (years) | 32.67 (±14.88) | 35.27 (±14.83) | 0.60 |
| Education (years) | 15.17 (±2.48) | 15.89 (±1.78) | 0.32 |
| Musical education (years) | 0.056 (±0.24) | 0 | 0.33 |
| Sex | 11F, 7M | 11F, 7M |  |
| Handedness | 2L, 16R | 1L, 17R | 0.55 |
| MBEA score | 21.55 (+/- 1.80)<br>Min: 16.83<br>Max: 24.17 | 26.60 (+/- 1.09)<br>Min: 24.5<br>Max: 28.5 | < 0.001 |
| MBEA pitch mean score | 20.67 (±2.08)<br>Min: 15.67<br>Max: 23.67 | 26.78 (±1.53)<br>Min: 23.33<br>Max: 29 | < 0.001 |
| PDT (semitones) | 1.72 (±1.46)<br>Min: 0.33<br>Max: 4.99 | 0.22 (±0.13)<br>Min: 0.08<br>Max: 0.54 | <0.001 |
| Musical emotion score (N=15 in both groups) | 3.85 (±0.87)<br>Min: 2.21<br>Max: 4.93 | 4.41 (±0.35)<br>Min: 3.86<br>Max: 5 | 0.032 |
| Speech perception score (N=15 in both groups) | 2.44 (±0.35)<br>Min: 1.86<br>Max: 3.14 | 2.13 (±0.30)<br>Min: 1.71<br>Max: 2.71 | 0.038 |

Prior to the study, 15 amusic and 15 control participants (among the cohort previously described) filled out a questionnaire about their relationships to music and language in their everyday life (based on questionnaires of McDonald & Stewart (2008)], Sloboda, Wise and Peretz (2005)], and Peretz et al. (2009)]). Among 90 questions, we selected 14 questions about musical emotions with 9 positive (e.g., “Music can relax me or calm me in a period of stress”) and 5 negative (“I never had a chill while listening to music”) statements. Participants rated their agreement to these statements from 1 (totally disagree) to 5 (totally agree). For each participant, an average musical emotions score was calculated (with a reverse coding for negative agreements). We also selected 7 questions about speech perception, excluding simple understanding of a language with 6 positive (e.g “Can you tell if someone has an accent?”) and one negative (“Do you have difficulties recognizing environmental sounds?”) questions. Participants rated to what extent these situations apply to them from 1 (almost every time) to 4 (almost never). For each participant, an average language comprehension score was calculated (with a reverse coding for the negative question; see **Table 1**). Both the musical emotions score (t(28) = 2.38, *p* = 0.032) and the language perception score (t(28) = 2.29, *p* = 0.038) differed significantly between the amusics and the controls, with amusics having a lower musical emotions score (i.e. a less emotional effect of music) and a higher language comprehension score (i.e. a less good language comprehension).

### 2.2. Stimuli

#### Sentences

Twenty sentences were selected from a larger material set of semantically neutral sentences uttered with different emotions by male and female actors. This selection was based on a pretest by a cohort of non-musician participants (N=16) in an explicit recognition task. The selected sentences were chosen according to their high recognition scores of the intended emotions (all superior to 80%, mean ± SD = 91.2 % ± 7.5). For each emotion (joy, neutral, sadness, anger, fear), four sentences were selected, half pronounced by a male voice and half by a female voice. For each emotion and gender, there were two sentences in French: “J’espère qu’il va m’appeler bientôt” (“I hope he will call me soon”), and “L’avion est presque plein” (“The plane is almost full”). Stimuli lasted on average 1470ms (+/- 278ms) and were equalized in Root Mean Square (RMS) amplitude.

#### Vowels

Twenty productions of the vowel /a/ were selected from a larger material set, all produced with female voices (Charpentier et al., 2018). All stimuli lasted 400ms and were equalized in RMS amplitude. This selection was based on a pretest by the same control participants as for the sentence material (N=16). Four stimuli for four basic emotions (joy, sadness, anger, fear) and neutrality were chosen according to their high recognition scores (all superior to 69%, mean ± SD = 82.9% ± 7.4).

Acoustic parameters (pitch mean, pitch slope, spectral flux mean, spectral flux slope, spectral spread mean, spectral spread slope, brightness mean, brightness slope, roughness mean, roughness slope, spectral centroid mean, spectral centroid slope, inharmonicity mean, inharmonicity slope, and attack time) of the stimuli were computed with the MIR toolbox (Lartillot & Toiviainen, 2007); **Figure 1**). Each parameter (except Attack Time) was computed with a frame of 50ms by default. We then computed the average of each parameters across time, and the slope of variation of these parameters across time. For sentences, the duration of the stimuli was also computed. For sentences and for vowels, an Analysis of Variance (ANOVA) was performed on each acoustic parameter with emotion (joy, sadness, anger, fear, neutral) as a between-stimuli factor. For sentences, there was a significant effect of emotion for pitch mean (F(4,15)=3.14, p=0.046), brightness mean (F(4,15) =3.2, p=0.044), spectral flux mean (F(4,15)=5.64, p=0.006), spectral flux slope (F(4,15)=6.47, p=0.003), spectral spread mean (F(4,15)=6.07, p=0.004), spectral spread slope (F(4,15)=6.62, p=0.003), roughness slope (F(4,15) =3.38, p=0.037), and spectral centroid mean (F(4,15)=4.14, p=0.019). According to Fisher LSD post-hoc tests, Joy had a significantly higher pitch than Neutrality (p=0.024). Joy was brighter compared to Neutrality (p=0.04). Joy and Fear had higher mean spectral flux than Sadness and Neutrality (all p<0.05). Anger had stronger slope of spectral flux than Sadness and Neutrality (all p<0.04). Joy, Fear and Anger had significantly lower spectral spread mean than Neutrality (all p<0.05). Anger had less steep spectral spread slope than Sadness and Neutrality (all p<0.02). Anger had significantly stronger roughness slope than Neutrality (p=0.04). Fear had significantly lower spectral centroid mean than Sadness (p=0.03).

For vowels, there was a significant effect of emotion for mean pitch (F(4,15) =23.24, p<0.001), mean brightness (F(4,15) =5.85, p=0.005), brightness slope (F(4,15)=4.32, p=0.016), mean spectral flux (F(4,15)=36.52, p<0.001), roughness slope (F(4,15)=3.2, p=0.043), spectral centroid slope (F(4,15)=6.29, p=0.004), and mean inharmonicity (F(4,15) =28.74, p <0.001). Fear had a significantly higher pitch than the other emotions (all p<0.01), Joy had a significantly higher pitch than Neutrality and Sadness (all p<0.004). Joy was significantly brighter than Anger, Sadness and Neutrality (all p<0.05), and Fear was significantly brighter than Sadness (p=0.05). Anger had a significantly less steep brightness slope than Joy and Neutrality (all p<0.039). Joy, Anger and Fear had significantly higher mean spectral flux than Sadness and Neutrality (all p<0.001), Sadness had a higher mean spectral flux than Neutrality (p=0.05). Fear had significantly stronger roughness slope than Sadness (p=0.037). Joy and Neutrality had significantly stronger spectral centroid slope than Sadness and Anger (all p<0.05). Joy and Anger had significantly higher inharmonicity means than Fear, Sadness and Neutrality (all p<0.01), Fear had a significantly higher inharmonicity mean than Neutrality (p=0.003).

**Figure 1:**
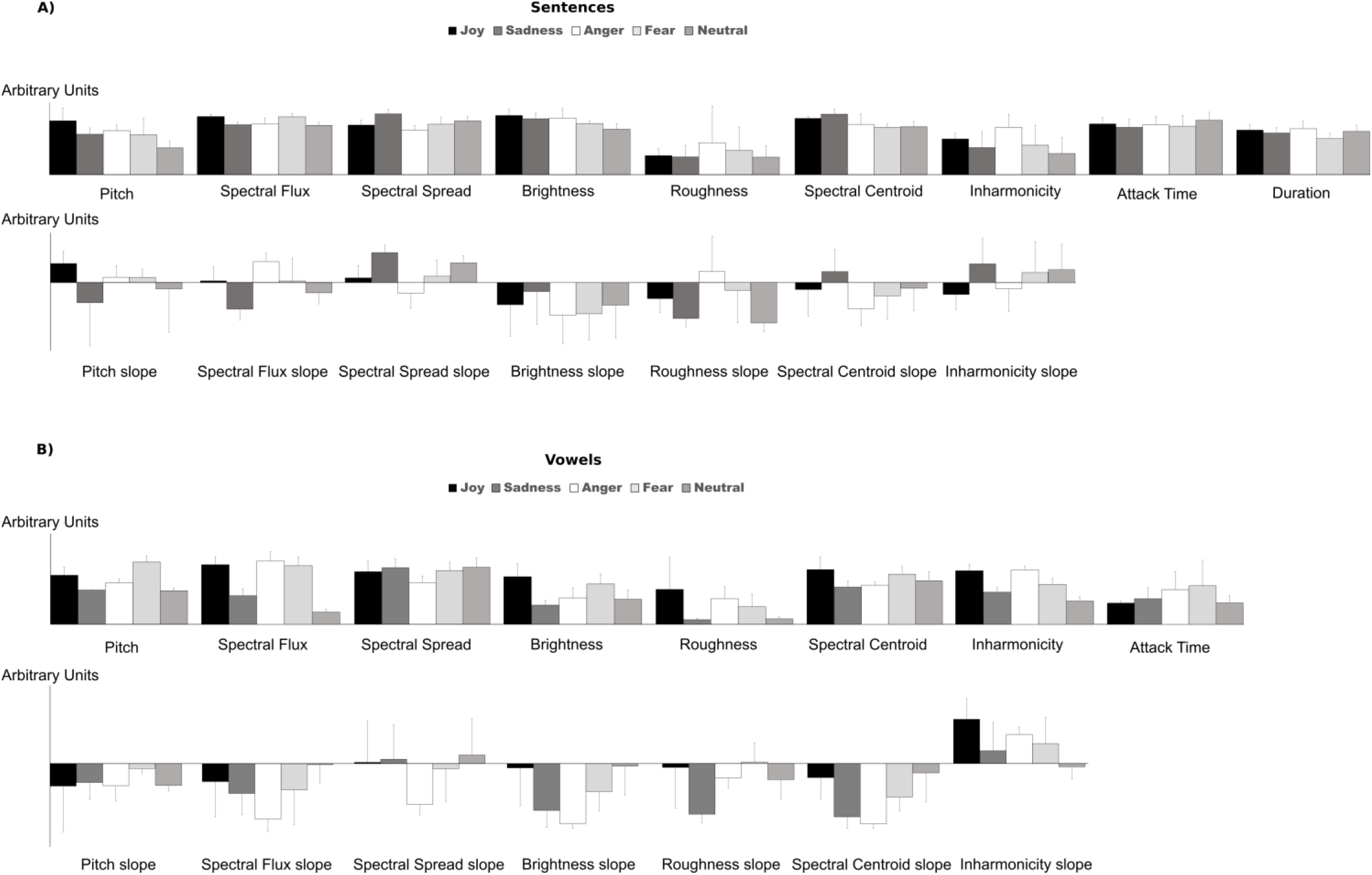
Acoustic parameters of the stimuli, averaged across all exemplars of a given emotion category. A)Parameters for the sentence stimuli. The first line represents the mean of the parameters (across time), the second line the slopes of their variation across time. There was a significant effect of emotion for pitch mean, spectral flux mean and slope, spectral spread mean and slope, brightness mean, roughness slope, and spectral centroid mean. B) Parameters for the vowel stimuli. There was a significant effect of emotion for pitch mean, spectral flux mean, brightness mean and slope, roughness slope, spectral centroid slope and inharmonicity mean. See main text for details. All units are arbitrary. Error bars correspond to the standard error of the mean. The slope sign refers to the general temporal ascending/descending tendency of the acoustical measure in stimuli.

#### 2.3. Procedure

The experiment took place in a sound-attenuated room. In each trial, participants listened to a stimulus and were then asked to select the recognized emotion from five options (joy, neutral, sadness, anger, fear). After having given their response, they were asked to rate the intensity of the selected emotion from 1 (not intense) to 5 (very intense), except for stimuli judged as neutral. After the intensity rating response, the following stimulus was automatically played after a variable delay of 1250 ms on average (ranging from 1000 to 1500 ms). The stimuli were presented in two blocks: sentences in one and vowels in another. The presentation order of the two blocks was counter-balanced across participants. Within a block, the presentation order of the stimuli was randomized for each participant, with the constraint that a given emotion cannot be presented more than three times in a row. Presentation software (Neurobehavioral systems, Albany, CA, USA) was used to present the stimuli to the participants and to record responses on a keyboard. The duration of the experiment was 15 minutes.

#### 2.4. Data analyses

For each participant and emotion, the percentage of correct responses was calculated, separately for sentences and vowels, and for correctly categorized trials, the average ratings of intensity were calculated, again separately for sentences and vowels. In light of recent criticism of classical frequentist analysis (Lee & Wagenmakers, 2014; Wagenmakers et al., 2018), a Bayesian approach was used to analyze the data. Bayesian analyses allow to perform model comparison and to select the best model (with the best evidence), given the data. We analyzed the data with the software JASP (Wagenmakers et al., 2017), with a 2×5 Bayesian mixed repeated-measures analysis of variance (ANOVA) on percentages of correct responses and intensity ratings with group (amusic or control) as between participants’ factor, and emotion (joy, sadness, fear, anger or neutral) as within-participant factor (note that for intensity rating, the emotion factor had only four levels as neutral stimuli were not rated for intensity). We also analyzed the data of sentences and vowels together with group (amusic or control) as between-participants factor, and emotion (joy, sadness, fear, anger or neutral, the latter only for percentage of correct responses) and material (sentences or vowels) as within-participant factors. We reported Bayes Factor (BF) as a relative measure of evidence. To interpret the strength of evidence, we considered a BF under three as weak evidence, a BF between three and 10 as positive evidence, a BF between 10 and 100 as strong evidence and a BF higher than 100 as a decisive evidence (according to (Lee & Wagenmakers, 2014). BF_10_ indicates the evidence of H1 compared to H0, BFinclusion indicates the evidence of one effect over all models. As no post-hoc tests with correction for multiple comparison have as yet been developed for Bayesian statistics (Wagenmakers et al., 2017), we used t-tests with Bonferroni correction for multiple comparisons.

In a second step, based on the percentage of correct responses of the emotion categorization task for the vowel material, we performed a multifactorial analysis (MFA) using the package FactoMineR (Lê et al., 2008) in R to analyze the underlying data structure (emotional space). We could not perform a MFA for sentence material due to high performance (ceiling effect). For each participant (18 amusics and 18 controls), we built a matrix with the responses for each stimulus in each of the five-emotion categories (20 in total), equivalent to a contingency table. Each participant and each stimulus were projected in the multi-dimensional space recovered by the analysis. R2 (correlation factor) is retrieved for each participant for each dimension and then compared between groups with t-tests. The acoustic parameters describing the stimuli were correlated with the coordinates of the stimuli in the multidimensional space (r coefficients are obtained for the projection of each acoustic parameter in each dimension) to uncover the relation between emotion dimensions and acoustic parameters.

## 3. Results

**Figure 2:**
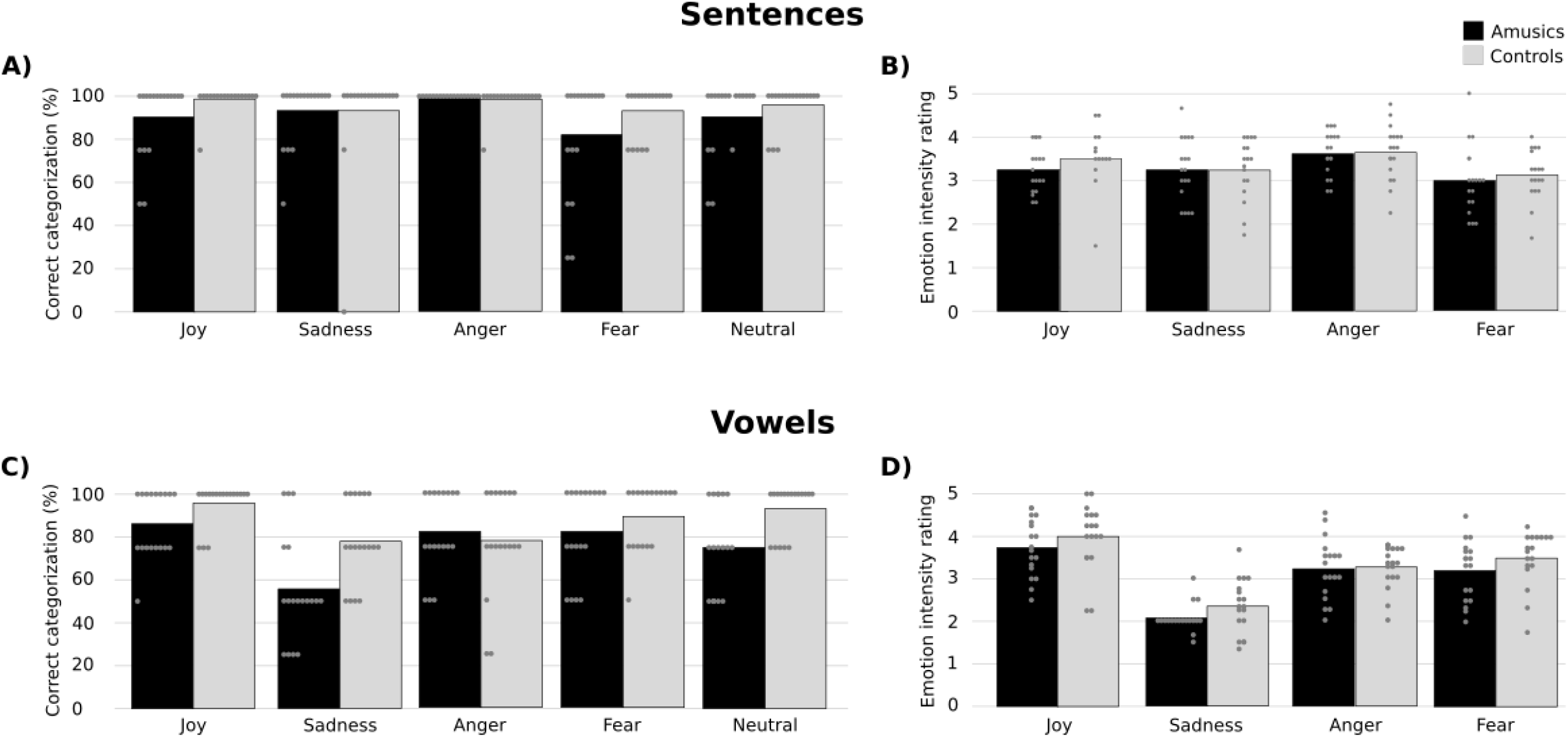
Percentage of correct categorization for amusic and control groups for sentence (A) and vowel (C) material. Mean intensity ratings for amusic and control groups, for sentence (B) and vowel (D) material. Bars represent the group means and dots correspond to individual data points. Amusics showed normal emotion recognition for sentences compared to controls. A ceiling effect was observed for sentences material as a majority of the participants were at 100*%*. Amusics showed a deficit for emotion recognition in vowels compared to controls. Some amusics were better than others (100*%* of correct answers) and performance were correlated with the PDT in amusic participants, with lower scores for participant with large PDTs. There were no significant group differences in emotion intensity ratings.

### 3.1. Sentences

#### *Emotion categorization* (Figure 2)

After comparison to the null model, the only model showing positive evidence was the one with only the main effect of emotion (BF_10_ = 5.39). All other models showed no significant evidence (BF_10_ < 3). This was confirmed by a positive specific effect of emotion only (BFinclusion = 3.85), all other specific effects were not significant (BF_inclusion_ < 0.5) (**Table 2**). According to t-tests with Bonferroni correction, Anger was significantly better recognized than Fear (t(34)= 3.55, pcorr<0.001).

**Table 2:** Results of the Bayesian mixed repeated measures ANOVAs on sentence stimuli, for correct categorization scores and intensity ratings. P(M): prior probability assigned to the model, P(M\data): probability of the model knowing the data, BF_M_: Bayesian Factor of the model, BF_10_: Bayesian Factor of the model compared to the null model.

|  | *Models* | *P(M)* | *P(M\|data)* | *BF<sub>M</sub>* | *BF<sub>10</sub>* | *error %* |
| --- | --- | --- | --- | --- | --- | --- |
| **Correct categorization** |  |  |  |  |  |  |
|  | Emotion | 0,2 | 0,525 | 4,416 | 5,39 | 1,345 |
|  | Emotion + Group | 0,2 | 0,274 | 1,509 | 2,813 | 1,403 |
|  | Null model (incl. subject) | 0,2 | 0,097 | 0,431 | 1 |  |
|  | Emotion + Group + Emotion × Group | 0,2 | 0,054 | 0,228 | 0,553 | 1,004 |
|  | Group | 0,2 | 0,05 | 0,211 | 0,515 | 1,261 |
| **Intensity** |  |  |  |  |  |  |
|  | Emotion | 0,2 | 0,679 | 8,479 | 3716,624 | 0,609 |
|  | Emotion + Group | 0,2 | 0,278 | 1,542 | 1521,765 | 0,803 |
|  | Emotion + Group + Emotion × Group | 0,2 | 0,042 | 0,176 | 230,261 | 2,025 |
|  | Null model (incl. subject) | 0,2 | 1,828e -4 | 7,314e -4 | 1 |  |
|  | Group | 0,2 | 6,943e -5 | 2,777e -4 | 0,38 | 0,632 |

Within each participant group (amusic or control), no significant Pearson-correlation was found between the categorization performance and the MBEA score (r(16)=0.004, p=0.99 for controls, r(16)=0.29, p=0.24 for amusics). Similarly, no correlation was found between categorization performance and the PDT (r(16)=-0.26, p=0.29 for controls, r(16)=-0.43, p=0.07 for amusics). However, a correlation was found between categorization performance and PDT over the two participant groups (r(34)=-0.43, p=0.009).

Confusion matrices are presented in **Table 3** for the two participant groups. These matrices showed similar patterns for amusic and control participants.

**Table 3:** Percent of answer types for each intended emotion for sentence material, averaged over the 18 participants for each group. Correct answers are on the diagonal.

|  | Answered |  |  |  |  |  |
| --- | --- | --- | --- | --- | --- | --- |
|  | Expected | *Joy* | *Sadness* | *Anger* | *Fear* | *Neutral* |
| Amusics |  |  |  |  |  |  |
|  | Joy | **90.28** | 2.78 | 0 | 6.94 | 0 |
|  | Sadness | 0 | **93.06** | 0 | 6.94 | 0 |
|  | Anger | 0 | 0 | **100** | 0 | 0 |
|  | Fear | 0 | 6.94 | 8.33 | **81.94** | 2.78 |
|  | Neutral | 0 | 8.33 | 1.39 | 0 | **90.28** |
| Controls |  |  |  |  |  |  |
|  | Joy | **98.61** | 0 | 0 | 1.39 | 0 |
|  | Sadness | 1.39 | **93.06** | 0 | 5.56 | 0 |
|  | Anger | 0 | 0 | **98.61** | 1.39 | 0 |
|  | Fear | 0 | 2.78 | 4.17 | **93.06** | 0 |
|  | Neutral | 0 | 2.78 | 1.39 | 0 | **95.83** |

#### Intensity ratings

The entire range (from 1 to 5) of intensity ratings was covered by the participants, showing that they fully used the subjective scale when rating the stimuli.

**Figure 3:**
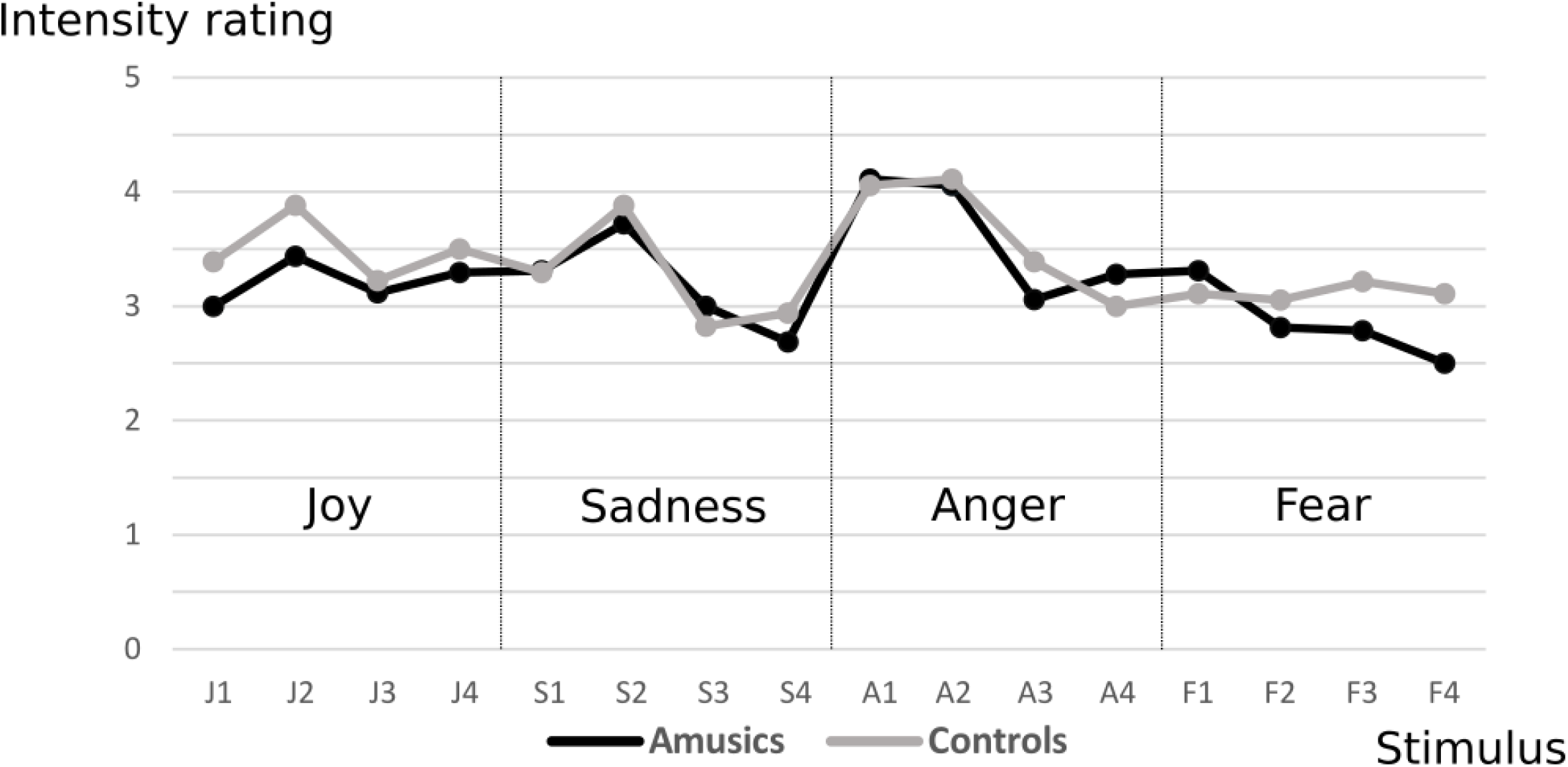
Means of emotion intensity ratings by stimulus for the amusic and the control group, for sentence material. Stimuli are classified by emotion. Ratings are given on a scale from 1 (not intense) to 5 (very intense). J1-4: Joy, S1-4: Sadness, A1-4: Anger, F1-4: Fear.

After comparison to the null model, the best model showing a decisive evidence was the one with the main effect of emotion only (BF10 = 3716.62) (**Figure 3**). This model was 2.44 times more likely than the model with the two main effects of group and emotion (BF10 = 1521.77) and 16.16 times more likely than the model with the two main effects of emotion and group and their interaction (BF10 = 230.26). The model with the main effect of group showed no evidence compared to the null model (BF10 =0.38). This was confirmed by a decisive specific effect of emotion only (BF_inclusion_ = 2642.28), all other specific effects were non-significant (BF_inclusion_ < 0.4) (**Table 2**). According to t-tests with Bonferroni correction, Fear was rated significantly lower than Joy and Anger (t(34)=3.41 pcorr=0.012 and t(34)=5.58 pcorr<0.001 respectively), and Sadness rated lower than Anger (t(35)=2.88 pcorr=0.042)^1^.

Amusic and control participants rated the stimuli similarly: amusics’ and controls’ intensity ratings correlated across stimuli (r(14)= 0.83, p<0.001)^2^. Moreover, both groups had similar reliability for intensity judgments (α Cronbach amusics=0.79 and α Cronbach controls =0.88, Fisher-Bonett Z=-1.08, p=0.14).

### 3.2 Vowels

#### *Emotion categorization* (Figure 2)

After comparison to the null model, the best model showing a decisive evidence was the one with the two main effects of group and emotion and their interaction (BF_10_ = 211784.75). This model was 2.44 times more likely than the model with the two main effects of group and emotion (BF_10_ = 86921.54) and 17.99 times more likely than the model with the main effect of emotion (BF_10_ = 11773.31). The model with the main effect of group showed positive evidence (BF_10_ =4.83). This was confirmed by a decisive specific effect of emotion (BFinclusion = 35508.55), a strong specific effect of group (BF_inclusion_ =16.91) and positive evidence of the interaction between emotion and group (BF_inclusion_ =8.58) (**Table 4**). According to t-tests with Bonferroni correction, Sadness was significantly less recognized than Joy, Fear, Anger and Neutrality (all pcorr<0.001). Amusics had lower recognition scores compared to controls only for Sadness (t(16)=2.76 pcorr=0.05) and for Neutrality (t(16)=3.42 pcorr=0.015) (other pcorr > 0.15).

**Table 4:**
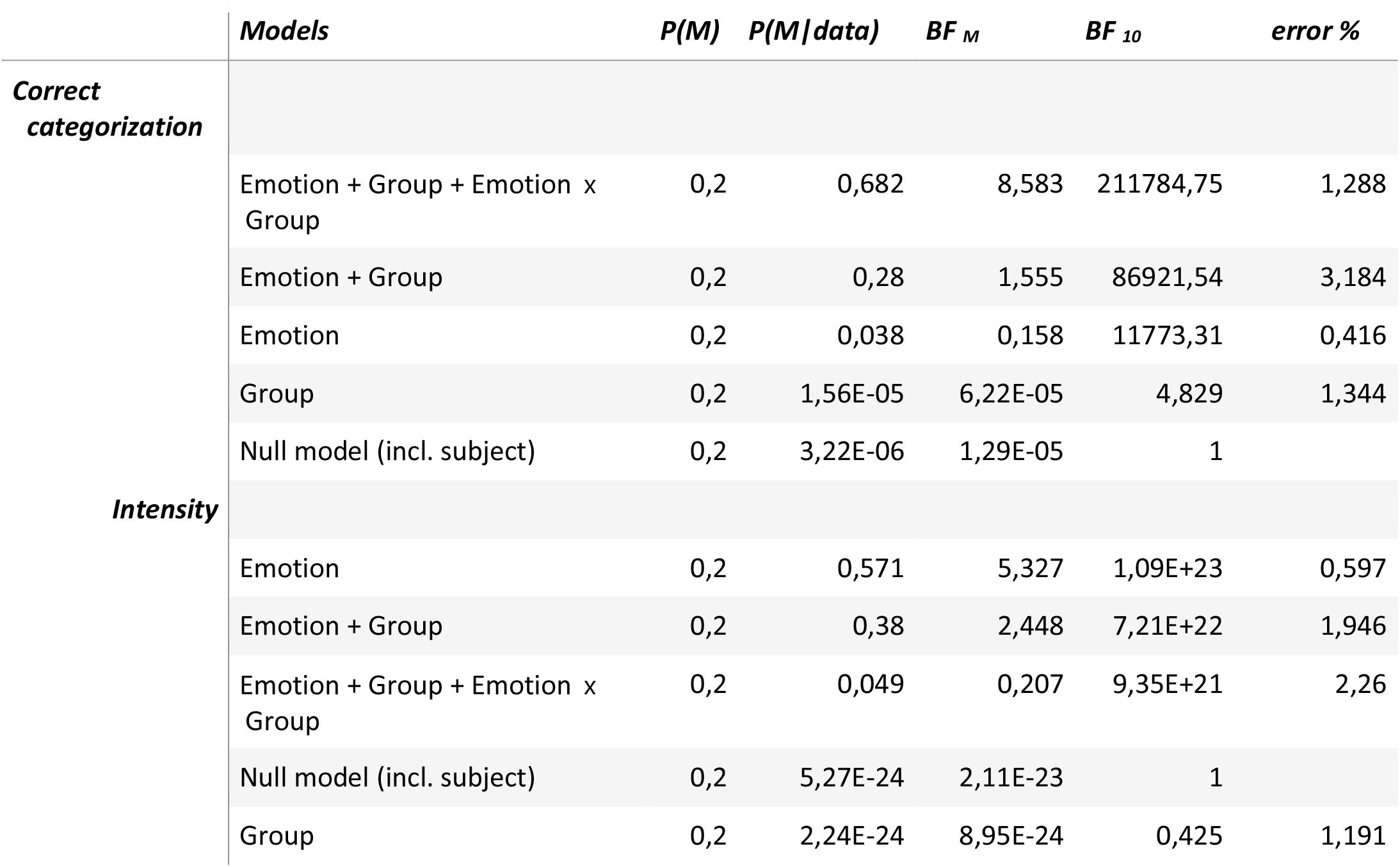
Results of the Bayesian mixed repeated measures ANOVAs on vowel stimuli, for correct categorization scores and intensity ratings. P(M): prior probability assigned to the model, P(M\data): probability of the model knowing the data, BF_M_: Bayesian Factor of the model, BF_10_: Bayesian Factor of the model compared to the null model.

Within each participant group (amusic or control), no significant Pearson-correlation was found between categorization performance and the MBEA score (r(16)=0.18, p=0.49 for controls, r(16)=0.26, p=0.29 for amusics). Similarly, no correlation was found between categorization performance and the PDT (r(16)=-0.070, p=0.78 for controls, r(16)=-0.32, p=0.2 for amusics). However, a correlation was found between categorization performance and the MBEA score over the two groups (r(34)=0.55, p<0.001). Similarly, a correlation was found between categorization performance and the PDT over the two groups (r(34)=-0.50, p=0.0021).

For each participant group, response confusions are shown in confusion matrices in **Table 5**. Interestingly, amusics seemed to show more confusion of sadness with other emotions, in particular with neutrality, than did controls. Amusics seemed to have a bias in the recognition of sadness in vowels towards neutrality. This point will be further explored in the MFA reported below.

**Table 5:** Percent of answer types for each intended emotion for vowel material, averaged over the 18 participants for each group. Correct answers are on the diagonal.

|  | Answered |  |  |  |  |  |
| --- | --- | --- | --- | --- | --- | --- |
|  | Expected | *Joy* | *Sadness* | *Anger* | *Fear* | *Neutral* |
| Amusics |  |  |  |  |  |  |
|  | Joy | **86.11** | 8.33 | 2.78 | 0 | 2.78 |
|  | Sadness | 5.56 | **55.56** | 0 | 6.94 | 31.94 |
|  | Anger | 2.78 | 0 | **81.94** | 6.94 | 8.33 |
|  | Fear | 6.94 | 6.94 | 2.78 | **81.94** | 1.39 |
|  | Neutral | 4.17 | 6.94 | 4.17 | 9.72 | **75** |
| Controls |  |  |  |  |  |  |
|  | Joy | **95.83** | 1.39 | 1.39 | 0 | 1.39 |
|  | Sadness | 1.39 | **77.78** | 1.39 | 1.39 | 18.06 |
|  | Anger | 1.39 | 5.56 | **77.78** | 8.33 | 6.94 |
|  | Fear | 5.56 | 5.56 | 0 | **88.89** | 0 |
|  | Neutral | 1.39 | 2.78 | 1.39 | 1.39 | **93.06** |

#### Intensity ratings

The entire range (from 1 to 5) of intensity ratings was covered by the participants, showing that they fully used the subjective scale to rate the stimuli.

**Figure 4:**
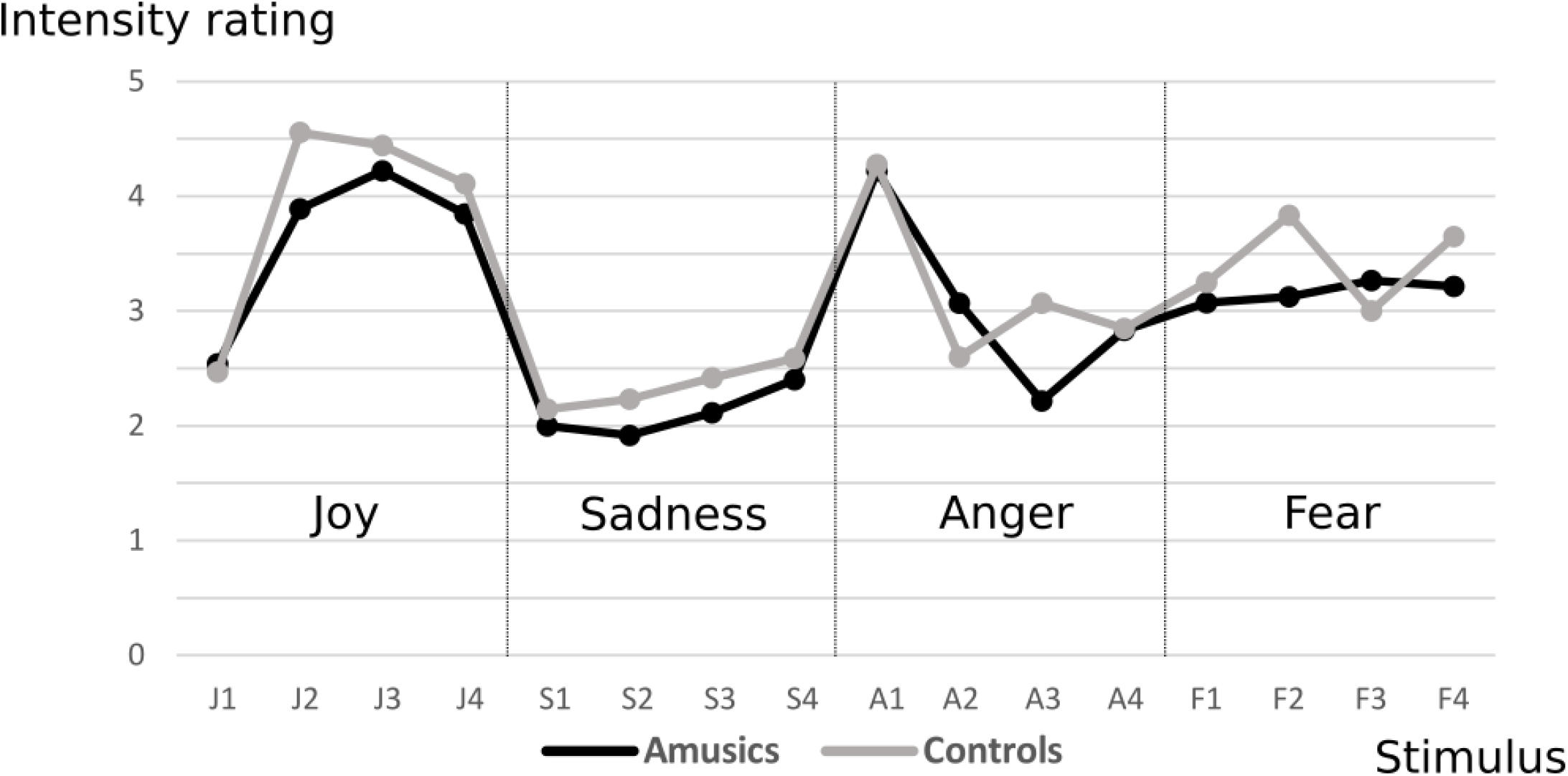
Means of emotion intensity ratings by stimulus for the amusic and the control groups, for vowel material. Stimuli are classified by emotion. Ratings are given on a scale from 1 (not intense) to 5 (very intense). J1-4: Joy, S1-4: Sadness, A1-4: Anger, F1-4: Fear.

After comparison to the null model, the best model showing decisive evidence was the one with the main effect of emotion only (BF_10_ = 1.09e+23) (**Figure 4**). This model was 1.51 times more likely than the model with the two main effects of group and emotion (BF_10_ = 7.22e+22) and 11.6 times more likely than the model with the two main effects of emotion and group and their interaction (BF_10_ = 9.35e+21). The model with the main effect of group showed no evidence compared to the null model (BF10 =0.43). This was confirmed by a decisive specific effect of emotion only (BFinclusion = 6e+15), all other specific effects were non-significant (BF_inclusion_ < 0.5) (**Table 4**). According to t-tests with Bonferroni correction, Sadness was rated significantly lower than Joy, Fear and Anger (all pcorr<0.001), and Anger and Fear rated lower than Joy (both pcorr<0.001)^3^.

Amusic and control participants rated the stimuli similarly (correlation between groups across stimuli: r(14)= 0.91, p<0.001)^4^. Moreover, both groups had similar reliability for intensity judgements (α Cronbach amusics=0.7 and α Cronbach controls =0.76, Fisher-Bonett Z=-0.43, p=0.33).

#### Multifactorial analysis

The multifactorial analysis revealed that four dimensions explained 69% percent of the variance in categorization data for vowels (**Figure A1**). The stimulus positions across these four dimensions confirmed the categorical structure of our stimulus set (**Figure 5A**). In particular, the first dimension corresponded to the separation of anger and joy stimuli, the second dimension to the separation of anger and neutral/sadness/fear stimuli, the third dimension to the separation of fear stimuli compared to the other emotions, and the fourth dimension corresponded to the separation of neutral and sadness stimuli. We correlated the acoustic parameters with the emotional space dimensions resulting from the multifactorial analysis (**Figure 5B**). Note in particular that the brightness mean (r(18)=-0.6, p=0.004) and slope (r(18)=-0.49, p=0.027), the spectral centroid mean (r(18)=-0.56, p=0.009) and slope (r(18)=-0.46, p=0.042) and the inharmonicity slope (r(18)=-0.46, p=0.043) of the stimuli correlated with the first dimension. The roughness mean (r(18)=0.56, p=0.011), the inharmonicity mean (r(18)=0.84, p<0.001) and slope (r(18)=0.48, p=0.032), the spectral flux mean (r(18)=0.8, p<0.001), and the spectral spread mean (r(18)=-0.58, p=0.007) and slope (r(18)=-0.47, p=0.038) of the stimuli correlated with the second dimension. The pitch mean (r(18)=0.82, p<0.001) of the stimuli correlated with the third dimension. Both the brightness slope (r(18)=0.49, p=0.029) and the spectral centroid slope (r(18)=0.56, p=0.01) of the stimuli correlated with the fourth dimension. This suggest that numerous acoustic cues are used to distinguish anger from other emotions in vowels: the major acoustic cue used to distinguish fear from other emotions was pitch, whereas the ones used to distinguish between neutral and sadness vowels were spectro-temporal cues, such as the slopes of brightness and spectral centroid. Interestingly, the third and fourth dimensions were the only dimensions to separate the two participant groups in terms of emotion classification (for dimension 3: cos2=0.259 and R2=0.81 (+/-0.043) for controls, cos2=0.231 and R2=0.67 (+/- 0.06) for amusics, and for dimension 4: cos2=0.150 and R2=0.62 (+/- 0.054) for controls, cos2=0.068 and R2=0.36 (+/-0.57) for amusics). The group comparison of individual R2 (**Figure 5C**) on the third and the fourth dimensions revealed a significant between-group difference (dimension 3: t(16)= 2.09, p=0.044, and dimension 4: t(16) = 2.87, p=0.007), with control participants contributing more strongly to these dimensions.

**Figure 5:**
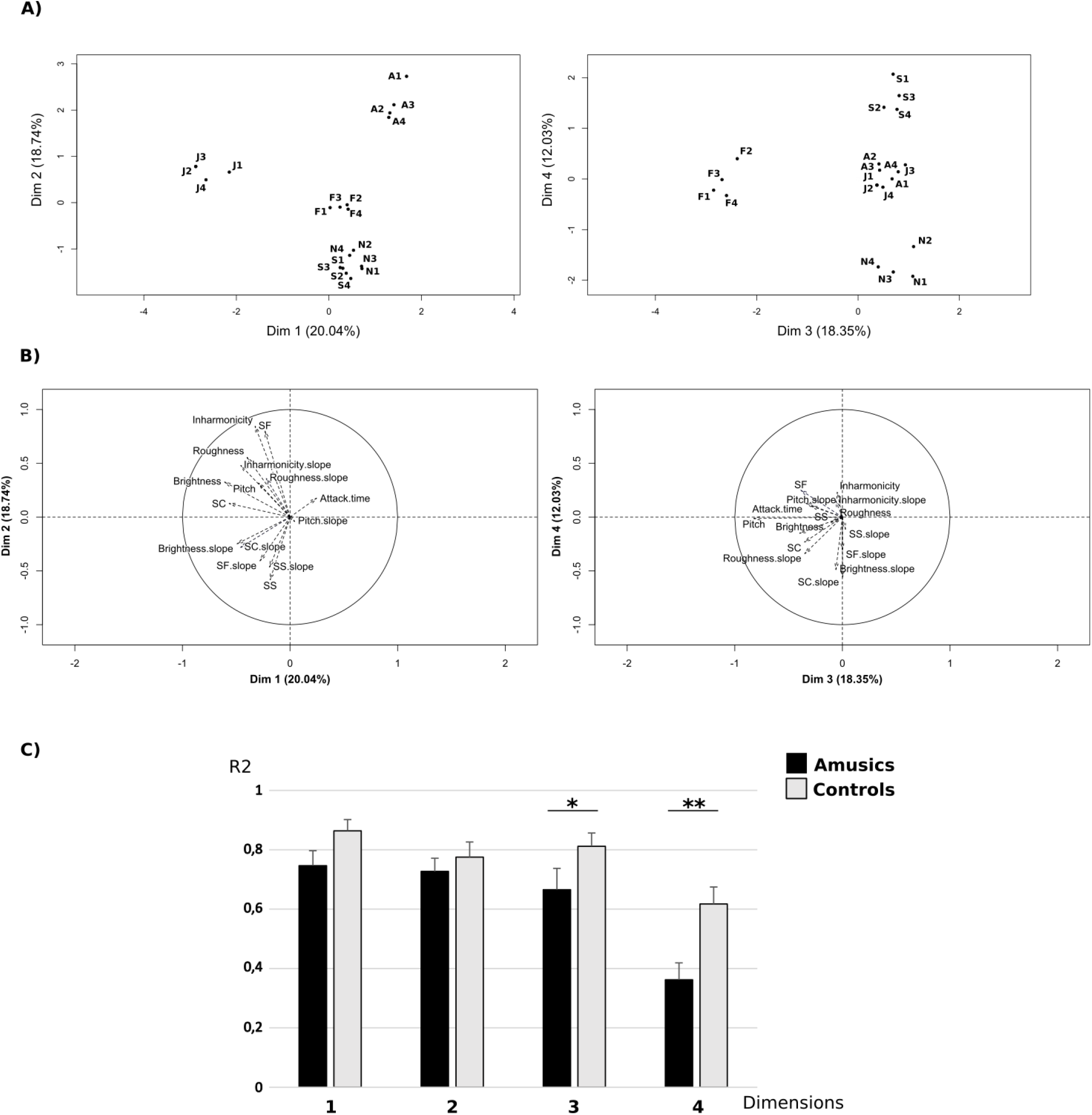
Results of the multifactorial analysis (MFA) on categorization data for vowel stimuli. Categorization data were explained by a four-dimensional model. The third and fourth dimensions differentiate between the two groups (amusics and controls). The third dimension corresponds to the separation of fear stimuli from other emotions. The fourth dimension corresponds to the separation of neutral and sadness stimuli. A) Representation of the stimuli in the four dimensions. B) Mean R2 correlation of the two groups (amusics and controls) for the first four dimensions of the MFA. The groups were mainly separated by the fourth dimension (p = 0.007), and to a lesser extent by the third dimension (p=0.044), error bars indicate standard error. C) Correlations between acoustic parameters describing the emotional stimuli with the four dimensions of the AFM. SF = Spectral Flux, SC = Spectral Centroid, SS = Spectral Spread. J1-4 = Joy, N1-4 = Neutral, S1-4 = Sadness, A1-4 = Anger, F1-4 = Fear.

### 3.3. Sentences and Vowels

#### Emotion categorization

After comparison to the null model, the best model showing a decisive evidence was the one with the three main effects of group, emotion, and material (vowels or sentences) and the interaction between material and emotion (BF_10_ = 1.02e+13). This model was 1.89 times more likely than the model with the three main effects of group, emotion and material and the interactions between material and group, and between material and emotion (BF_10_ = 5.41e+12). The best model was also 2.03 times more likely than the model with the three main effects of group, emotion and material and the interactions between material and emotion, and between group and emotion (BF_10_=5.02e+12). The best model was 3.25 times more likely than the model with the three main effects of group, emotion and material and the interactions between material and group, between material and emotion, and between group and emotion (BF_10_=3.14e+12). The best model was 4.81 times more likely than the model with the three main effects of group, emotion and material and the interactions between material and group, between material and emotion, between group and emotion and between material, group and emotion (BF_10_=2.12e+12). Finally, the best model was 5.34 times more likely than the model with the two main effects of emotion and material and the interaction between the two (BF_10_=1.91e+12). All other models were more than 2513 times less likely than the best model (all BF_10_<4e+9). This was confirmed by a decisive evidence for the specific effects of material (BF_inclusion_ =7.44e+10), emotion (BF_inclusion_ = 97881.46), and their interaction (BF_inclusion_ =5818.48), and a positive evidence for the specific effect of group (BF_inclusion_ = 4.83). See supplementary data for the full result table (**Table A1**). Amusics had lower scores than Controls and Vowels were less well recognized than Sentences. According to t-tests with Bonferroni correction, Sadness was significantly less well recognized than Joy and Anger (t(71)=4.06 pcorr<0.001 and t(71)=3.05 pcorr=0.03 respectively). Sadness (t(35)=4.76 pcorr<0.001) and Anger (t(35)=5.39 pcorr<0.001) were less well recognized in Vowels than in Sentences.

#### Intensity ratings

After comparison to the null model, the best model showing a decisive evidence was the one with the two main effects of emotion and material and the interaction between the two (BF_10_ = 3.92e+29). This model was 2.05 times more likely than the model with the three main effects of group, emotion and material and the interaction between material and emotion (BF_10_ = 1.91e+29). All other models were more than 10 times less likely than the best model (all BF_10_<4e+28). This was confirmed by decisive evidence for the specific effects of material (BF_inclusion_ =∞), emotion (BF_inclusion_ =∞), and their interaction (BF_inclusion_ =9.76e+15), no evidence for other main effects or interaction (all BF_inclusion_ <0.22). See supplementary data for full table (**Table A1**). According to t-tests with Bonferroni correction, Joy was rated with stronger intensity than Fear and Sadness (t(71)=5.91 pcorr<0.001) and t(71)=7.46 pcorr<0.001 respectively), Anger rated as more intense than Sadness (t(71)=7.65 pcorr<0.001), Fear rated higher than Sadness (t(71)=4.21 pcorr<0.001). Joy was rated as more intense in vowels than sentences (t(35)=3.78 pcorr<0.001), and Sadness and Anger rated higher in sentences than vowels (t(35)=9.42 pcorr<0.001 and t(35)=3.37 pcorr=0.01 respectively).

## 4. Discussion

The aim of the present study was to investigate emotional prosody perception in congenital amusia. To do so, we used sentences with neutral semantic content and short vowels (/a/) to investigate the explicit recognition of emotions and related intensity ratings. The stimuli were pronounced to express four different basic emotions: joy, fear, anger, and sadness, and with a neutral voice. To further investigate the perception of emotions in speech, and based on recent findings about congenital amusia and music-evoked emotions (Lévêque et al., 2018), participants were also asked to rate the intensity of the emotion previously recognized. Overall, the results revealed an explicit deficit in the amusic group for emotion recognition in vowels, but not in sentences. This deficit was most pronounced for sadness and neutrality, which tended to be confounded with each other or with fear more often by amusics than by controls for the vowel material. These impairments in emotional prosody recognition seem to be linked to a deficit in the processing of pitch and spectro-temporal parameters of the stimuli, with a relative preservation of harmonicity and roughness processing in congenital amusia. Despite this explicit deficit, implicit judgments on these emotions seemed to be preserved, as intensity ratings were similar in both participant groups for both materials.

### 4.1. Deficit of emotional prosody processing in congenital amusia

For the sentence material, amusics’ performance did not differ from controls’ performance: both groups were equally able to recognize the expressed emotions. This was confirmed by the similar confusion matrices between the two participant groups. According to our acoustic analysis on the sentence material, we found significant differences between emotions for a variety of pitch-related cues, spectral cues and spectro-temporal cues (slopes of spectral parameters variation across time). We suggest that these numerous acoustic cues associated with the rather long duration of our stimuli (an average duration of 1470 ms per sentence) were sufficient for participants of the two groups to distinguish emotions. The combination (and accumulation over time) of these acoustic cues could explain why amusics’ emotion recognition was similar to that of controls, as previously shown by Lolli et al. (2015).

The results for the full-sentence stimuli showed no evidence for an impairment in emotion recognition in amusia. Note that near-ceiling performance was observed in both participant groups. To further investigate possible group differences, we also presented participants with a simpler 400 ms vowel material where temporal cues are minimal, and less acoustic information is available than in full sentences. For the vowel material, we observed a group effect, with lower performance in the amusics relative to the controls. Interestingly, this was accompanied by an interaction between group and emotion. Specifically, amusics had more difficulties recognizing sadness and neutrality than did controls. These results were confirmed by the confusion matrices of each group, showing more confusion between sadness and neutrality in the amusic group than in the control group (these two emotions being also sometimes misclassified as fear). In controls, this bias was less marked, and controls had more difficulties recognizing anger than sadness, which was not the case for amusics. A multifactorial analysis confirmed that all stimuli were well separated across the five emotions. This analysis also revealed how acoustic parameters determined emotion categorization. Correlations between acoustic parameters and the fourth axis of the emotional space (recovered by the multifactorial analysis) separating sadness and neutral stimuli revealed the similarity of several acoustic features between neutral and sadness stimuli, the most frequent confusion in the amusic group. Significant correlations, meaning that the acoustic feature could theoretically be used to distinguish the two emotions, occurred only for (1) brightness slope and (2) spectral centroid slope. These spectro-temporal cues have previously been shown to be perceptually less salient than other acoustic cues in non-amusic, normal-hearing listeners (Caclin, McAdams, Smith, & Winsberg, 2005), showing that neutral and sadness stimuli are perceptually rather close. This suggests that amusic individuals have difficulty detecting contrasts between emotions with less salient variations of acoustic parameters. Indeed, the differences between neutrality and sadness are minimal in pitch, brightness, roughness, and inharmonicity, which are often used by non-amusic participants to recognize an emotion (Marin et al., 2015).

The first two dimensions of the multifactorial analysis correlated with several acoustic cues, such as roughness and inharmonicity. No deficit was observed in the amusic group on these two dimensions. This confirmed previous results showing that amusic individuals judge affective sounds mostly based on roughness and sometimes on harmonicity, whereas, relative to controls, they have reduced sensitivity to other acoustic parameters (Marin et al., 2015). The third dimension of the multifactorial analysis correlated with pitch, and the results of the amusic and control groups differ significantly for this dimension, confirming the amusic deficit in emotional prosody recognition when pitch is needed (Liu et al., 2015; Marin et al., 2015). The findings suggest that amusics need more variation of acoustic cues in speech to understand emotional prosody and rely more on accumulation of acoustic cues over time to determine the emotion.

Our additional analysis for sentence and vowel materials revealed that amusics had lower recognition scores overall. We observed a significant interaction between emotion and material showing that, depending on the material (sentences or vowels), participants did not recognize all emotions in the same way. Post hoc tests showed that sadness was less well recognized in vowels than in sentences, possibly revealing the need to process subtler acoustic cues and the less dynamic cues in sad vowel stimuli compared to sentences. The second best model of our Bayesian analysis included the interaction of group and material, showing a larger deficit of amusics with vowel material relative to other group/material combinations. Even if this interaction was not present in the best model, the second model which included this interaction was only 1.89 times less likely than the best model. This overall Bayesian analysis thus confirmed potential differences between the processing of sentence and vowel materials and revealed specific deficits of amusics with vowels, but not sentences.

### 4.2. Preserved implicit processing of emotional prosody in congenital amusia

To determine whether congenital amusic individuals were also impaired for implicit processing of emotional prosody, participants were asked to provide intensity ratings to each perceived emotion. Previous studies with patients with acquired amusia revealed a possible dissociation of emotion processing between recognition and the intensity ratings (Hirel et al., 2014; Peretz, Gagnon, & Bouchard, 1998). These studies showed a spared recognition of musical emotions whereas intensity ratings were impaired. Interestingly, here we observed unimpaired performance for the intensity ratings in congenital amusia, both for sentence and vowel stimuli. Indeed, we observed similar intensity ratings in both participant groups, with ratings differing only between emotions. Furthermore, the analysis on intensity ratings with sentences and vowels together revealed only two main effects of emotion and material, along with no group effect or interaction. This means that amusic participants’ intensity ratings of the stimuli were similar to those of the controls, regardless of the material used. Our analysis showed clear differences between intensity ratings for the different emotions, which was more pronounced in vowels than in sentences. These observations suggest that amusic participants were able to rate the intensity of the different emotions, even if they did not recognize them as well as did the controls. Results obtained with intensity judgements on all stimuli, including those not correctly categorized, showed the same pattern of results with a main effect of emotion. A small effect of group was revealed in the vowel intensity analysis, but the BFinclusion of the group factor was very small (<3), revealing a non-significant effect of group in the model and a stronger effect of emotion. Moreover, the correlation of intensity ratings across the stimuli revealed that both groups rated the stimuli the same way across emotions.

These analyses confirmed that emotion categorization and intensity ratings can be at least partially separated as two processes within global emotion processing as has been previously described in patients with acquired amusia or musical anhedonia (Hirel et al., 2014; Peretz et al., 1998), and that the deficit in amusia occurs mainly in explicit emotion categorization, whereas implicit emotional prosody processing is relatively preserved. Previous results support this conclusion: in a recent study with amusic participants performing explicit tasks on musical material, the results suggested explicit tonal processing difficulties alongside preserved implicit processing (Tillmann, Lalitte, et al., 2016). Indeed, when explicitly asked to judge the tonality of musical material, amusic participants had difficulties, but when the judgement was implicit, they performed as well as did controls (Tillmann, Lalitte, et al., 2016; Zendel et al., 2015). Similarly, our findings on verbal material showed some impairment of explicit recognition and categorization of emotion in vowels, but preserved implicit knowledge for the processing of the intensity of the emotional content in amusia.

In sum, previous findings have suggested preserved implicit processing capacities in congenital amusia for musical materials (Lévêque et al., 2018; Marin et al., 2015; Moreau et al., 2009; Omigie et al., 2013; Peretz et al., 2009; Tillmann et al., 2014, 2012; Tillmann, Lalitte, et al., 2016), and the present study expands these findings to a new domain, suggesting preserved implicit processing of pitch and spectro-temporal auditory features for verbal materials as well. This preserved implicit processing could provide a basis on which to develop new methods for auditory rehabilitation in amusia.

### 4.3. Understanding the deficit of congenital amusia beyond impaired pitch processing in musical perception

Various acoustic cues differentiate the emotions of the stimuli in the present study. For the vowels, the variations in acoustic parameters could be linked with the differences in recognition accuracy between groups. In the multifactorial analysis, pitch was correlated with the third dimension, which was where the amusic group’s performance differed significantly different from that of the controls. This provided support for the hypothesis of a pitch perception deficit in amusia (Liu et al., 2015; Marin et al., 2015). In addition, the fourth dimension of the multifactorial analysis revealed a significant difference between amusics and controls and correlated with spectro-temporal variations of the stimuli, namely the slopes of the acoustic parameters, in particular brightness and spectral centroid. This observation suggests that amusics could have a subtle deficit in the perception of these acoustic parameters, leading to a deficit in perception of a particular set of aurally-presented emotions. This deficit in spectro-temporal variation processing in amusia leads us to re-evaluate the potential pitch-perception deficit. Indeed, several studies have highlighted a contour-related pitch-perception deficit in amusia (Foxton, Dean, Gee, Peretz, & Griffiths, 2004; Patel et al., 2005). In agreement with these findings, the results of the multifactorial analysis seem to suggest a deficit in the processing of dynamic spectral information in congenital amusia. This is also in keeping with the deficit in short-term memory for pitch sequences which characterize amusic participants (Tillmann, Lévêque, et al., 2016). Finally, comparing results from perception of emotion in both sentences and vowels also confirmed that when speech prosody stimuli are long enough, accumulation of evidence allows the amusics to perform equally as well as controls for recognition of emotions (Lolli et al., 2015).

Moreover, our results lead to a better understanding of pitch processing in congenital amusia beyond musical material. Indeed, congenital amusia was first described as a music processing impairment (Ayotte et al., 2002; Peretz et al., 2003). However, recently, evidence has begun to accumulate that indicates that the pitch deficit in amusia extends to verbal materials as well (Nguyen et al., 2009; Tillmann, Burnham, et al., 2011; Tillmann, Rusconi, et al., 2011). This means that congenital amusia is a deficit not only of music perception but also of speech processing, in particular emotional and intentional prosody as well as tonal language, when pitch carries relevant information (Lolli et al., 2015; Nguyen et al., 2009; Patel, Wong, Foxton, Lochy, & Peretz, 2008; Thompson et al., 2012). Even though this deficit is small, here only observed with vowels, we also observed a difference in the experience of music and speech of amusics in their everyday life, compared to controls. Similar to what has been shown before, the musical emotion questions in the questionnaire revealed that amusic individuals less frequently experience emotions when listening to music (Mcdonald & Stewart, 2008; Omigie et al., 2012). More interestingly, the speech portion of the questionnaire showed a deficit in amusia for processing speech in a context of subtle acoustic variations (such as accent, intentional prosody). Even if this questionnaire was not specific for emotional prosody, it did reveal a small deficit for speech in amusia, beyond music, that could impact everyday life. For example, this impairment could become problematic when speech is presented in degraded conditions, such as speech in noise, speech in speech, or speech over the phone. These conditions of hearing have been shown to challenge comprehension in normal-hearing participants (Oxenham, 2008, 2012), particularly for the elderly, and could be even more difficult for amusic participants.

## 5. Conclusion

Overall, the results of the present study allow for a better understanding of the perceptual deficits of congenital amusia regarding speech material. They revealed a deficit in emotional prosody processing in amusia by examining emotion recognition of sentences and short vowels. The results suggest that the use of acoustic-constrained material can reveal subtle deficits in congenital amusia and allow for a better definition of acoustic cues that are critical for emotion perception in speech and its impairment in amusic individuals. The results also demonstrate a dissociation between explicit and implicit processing of emotions in congenital amusia (Lévêque et al., 2018). This finding contributes to a better understanding of the complex relationship between music and emotional prosody processing, and provides further elements for the comprehension of fine acoustic structures underlying music and speech appraisal.

## Acknowledgments

This work was supported by a grant from the Research National Agency (ANR-11-BSH2-001-01 to B. Tillmann and A. Caclin. This work was conducted in the framework of the LabEx CeLyA (‘‘Centre Lyonnais d’Acoustique”, ANR-10-LABX-0060) and of the LabEx Cortex (‘‘Construction, Function and Cognitive Function and Rehabilitation of the Cortex”, ANR-11-LABX-0042) of Université de Lyon, within the program ‘‘Investissements d’avenir” (ANR-11-IDEX-0007) operated by the French National Research Agency (ANR). This work was realized as part of a Marie Curie fellowship (International Incoming Fellowship) of A.Bhatara. We would like to thank Céline Tourlonnias for pretests with the sentence material. We also thank Lison Fanuel for her help with the Bayesian Analysis.

## Footnotes

1 We run another Bayesian ANOVA with intensity ratingsof all the trials (correct or not). This confirmed the decisive evidence of the best model with the maineffect of emotion (BF_10_ =2.68.106). This model was 2.25 times more likely than the model with the two main effects of group and emotion (BF_10_ = 1.19.106) and 12.18 times more likely than the model with the two main effects of emotion and group and their interaction (BF_10_ = 223230). This was confirmed by a decisive specific effect of emotion only (BF_inclusion_ = 1.95e+6), all other specific effects were non-significant (BF_inclusion_ < 0.35). T tests with Bonferroni correction confirmed that Anger was rated higher than Sadness, Fear and Joy (all pcorr<0.03), and Joy rated higher than Fear (pcorr<0.001).

2 We run another correlation measure with intensity ratings for all the trials (correct or not). It confirms that amusic and control participants rated the stimuli similarly: amusics’ and control’s intensity ratings correlated across stimuli (r(14)=0.84, p<0.001).

3 We run another Bayesian ANOVA with intensity ratings of all the trials (correct or not). This confirmed the decisive evidence of the best model with the main effect of emotion (BF_10_ =2.68.10^6^). This model was 2.25 times more likely than the model with the two main effects of group and emotion (BF_10_ = 1.19.10^6^) and 12.18 times more likely than the model with the two main effects of emotion and group and their interaction (BF_10_ = 223230). This was confirmed by a decisive specific effect of emotion only (BF_inclusion_ = 1.95e+6), all other specific effects were non-significant (BF_inclusion_ < 0.35). T tests with Bonferroni correction confirmed that Anger was rated higher than Sadness, Fear and Joy (all pcorr<0.03), and Joy rated higher than Fear (pcorr<0.001).

4 We run another correlation measure with intensity ratings for all the trials (correct or not). It confirms that amusic and control participants rated the stimuli similarly: amusics’ and control’s intensity ratings correlated across stimuli (r(14)=0.84, p<0.001).

## Abbreviations

MBEA: Montreal Battery for the Evaluation of Amusia
PDT: Pitch Discrimination Threshold

